# Endothelial Nitric Oxide Synthase Regulates Lymphatic Valve Specification By Controlling β - catenin Signaling During Embryogenesis

**DOI:** 10.1101/2023.04.10.536303

**Authors:** Drishya Iyer, Diandra Mastrogiacomo, Kunyu Li, Richa Banerjee, Ying Yang, Joshua P. Scallan

## Abstract

**Objective:** Lymphatic valves play a critical role in ensuring unidirectional lymph transport. Loss of lymphatic valves or dysfunctional valves are associated with several diseases including lymphedema, lymphatic malformations, obesity, and ileitis. Lymphatic valves first develop during embryogenesis in response to mechanotransduction signaling pathways triggered by oscillatory lymph flow. In blood vessels, eNOS (gene name: *Nos3*) is a well characterized shear stress signaling effector, but its role in lymphatic valve development remains unexplored.

**Approach and Results:** We used global *Nos3^−/−^* mice and cultured hdLECs to investigate the role of eNOS in lymphatic valve development, which requires oscillatory shear stress signaling. Our data reveal a 45% reduction in lymphatic valve specification cell clusters and that loss of eNOS protein inhibited activation of β-catenin and its nuclear translocation. Genetic knockout or knockdown of eNOS led to downregulation of β-catenin target proteins *in vivo* and *in vitro*. However, pharmacological inhibition of NO production did not reproduce these effects. Coimmunoprecipitation experiments reveal that eNOS forms a complex with β-catenin and their association is enhanced by oscillatory shear stress. Finally, genetic ablation of the Foxo1 gene enhanced FOXC2 expression and rescued the loss of valve specification in the eNOS knockouts.

**Conclusion:** In conclusion, we demonstrate a novel, nitric oxide-independent role for eNOS in regulating lymphatic valve specification and propose a mechanism by which eNOS forms a complex with β-catenin to regulate its nuclear translocation and thereby transcriptional activity.

## INTRODUCTION

Lymphatic vessels are required to maintain tissue fluid homeostasis by absorbing interstitial fluid and delivering it to the blood vessels^1^. Lymph transport against an increasing pressure gradient is facilitated by the growth of intraluminal, bicuspid valves during embryonic development that prevent lymph backflow^2^. Several diseases are associated with the loss of lymphatic valves or dysfunctional valves, including congenital lymphedema, cancer-associated lymphedema, obesity, ileitis, and lymphatic malformations^3–8^.

Lymphatic valve development in the mesentery occurs in response to the onset of lymph flow at embryonic day E15.5^9^. Lymph flow reversal at the branchpoints of the mesenteric lymphatic network generates an oscillatory shear stress (OSS) on the surface of the lymphatic endothelial cells (LECs)^9, 10^. In response to OSS, the transcription factor β-catenin translocates to the nucleus and upregulates the genes, PROX1 and FOXC2, in subsets of LECs, marking the specification of LEC clusters that later form valves^11^.

Mechanotransduction signaling in the lymphatic vasculature was recently shown to lead to AKT phosphorylation in a VE-cadherin-dependent manner and the AKT signaling pathway was partially responsible for lymphatic valve development^12^. A major target of AKT in the lymphatic valve leaflets is FOXO1, which is a repressor of lymphatic valve formation^13^. However, in blood vessels, it is well established that mechanotransduction signaling leads to phosphorylation of AKT and the S1177 residue of endothelial nitric oxide synthase (eNOS)^14^. While a previous study showed that eNOS expression is induced shortly after the onset of lymph flow and before the appearance of PROX1^high^ valve specification clusters in the mouse mesentery^9^, it remains unclear whether and how eNOS signaling regulates lymphatic valve development. Here, we show that eNOS plays an early role in lymphatic valve specification by regulating the transcription factor β-catenin and elucidate a mechanism by which eNOS forms a complex with β-catenin to regulate its transcriptional activity.

## METHODS

### Mice

The *Nos3^−/−^* and *Foxo1^flox^* strains were obtained from the Jackson Laboratory. The *Prox1-GFP, Ctnnb1^flox^, Prox1CreER^T2^* (TMak), and *Ctnnb1^ex3(loxP)^* strains were described previously^15–18^. All mice were maintained on a pure C57BL/6J background and both sexes were used. Tamoxifen (TM) was dissolved in sunflower oil and ethanol for a final stock of 20mg/mL with 5% ethanol. Embryonic deletion of Foxo1 and exon 3 of β-catenin was induced by intraperitoneal injection of 5mg TM into pregnant dams on E14.5. Postnatal deletion of β-catenin was induced by subcutaneous injection of 100ug of TM at P1 and P3. Postnatal recombination of exon 3 of β-catenin was induced by subcutaneous injection of 100ug of TM once on P0.

### Generation of Prox1CreERT2 Strain

Difficulty breeding a previous *Prox1CreER^T2^* with *Ctnnb1^flox^* to homozygosity indicated that these two alleles resided on the same chromosome. Therefore, we generated a new *Prox1CreER^T2^* strain by inserting a CreER^T2^ followed by a SV40 polyA signal right after the start codon in the RP23-360I16 BAC clone. Random transgenesis generated 4 founders and the highest efficiency founder was selected for propagation. With repeated breeding, homozygous *Prox1CreER^T2^;Ctnnb1^flox/flox^* mice were eventually obtained with both strains, presumably due to crossover events.

### Quantification of valve specification clusters, lymphatic valves and branchpoints

Mesenteries from embryos or postnatal pups were harvested along with the intestine and pinned onto a Sylgard 170 cushion. After immunostaining with PROX1 and FOXC2, the mesenteries were imaged on an inverted fluorescence microscope (Axio Observer Z1, Zeiss). Alternatively, tissues expressing the Prox1GFP reporter were imaged under a Zeiss Axio Zoom V16 fluorescence stereoscope. PROX1^hi^ cell clusters (cell clusters having 10 or more cells expressing high levels of PROX1) or PROX1^hi^ valves were quantified with Fiji (NIH ImageJ) software, which was also used to measure length of the large collecting vessels and the thinner pre-collecting vessels located near the intestinal wall. Between 4-5 arcades, defined as a large collecting vessel with its associated pre-collecting vessels, were imaged in each mesentery for later valve analysis.

### Quantification of PROX1 and FOXC2 expression *in vivo*

Mesenteries from P3 pups were collected and immunostained with PROX1, FOXC2 and VE-cadherin. High magnification confocal images of lymphangion and valve areas were obtained on a Leica TCS SP8 laser scanning confocal microscope. Using Fiji software, a ROI was drawn around each lymphangion or valve and added to ‘ROI manager’. The area of each ROI was the same for all lymphangion and valve regions. The mean pixel intensity was measured in each ROI in each image. Between 4-6 lymphangion or valve areas were imaged and analyzed from 5-6 control and knockout littermates.

### Whole-mount immunostaining procedure

All steps were carried at out 4µC on an orbital shaker (Belly Dancer, IBI Scientific) unless otherwise mentioned. The mesentery along with the intestine was harvested and pinned onto a Sylgard 170 cushion and subsequently fixed overnight in 1% paraformaldehyde. The tissues were washed the next day with PBS three times 10 minutes each after which the whole-mount immunostaining procedure was carried out in the following steps: (1) Permeabilize the tissues with PBS + 0.3% Triton X-100 (PBST) for one hour; (2) Block the tissues with 3% donkey serum in PBST for two hours; (3) Incubate the tissues with primary antibodies dissolved in PBST overnight; (4) Wash the tissues with PBST five times 15 minutes each; (5) Incubate the tissues with secondary antibodies dissolved in PBST for 2 hours at room temperature; (6) Wash the tissues with PBST five times 15 minutes each; (7) Mount tissues on glass slides (Superfrost Plus Microscope slides, Fisherbrand) using ProLong Diamond Antifade Mountant containing DAPI (Invitrogen) and store slides at 4µC overnight. Tissues were imaged on a fluorescence microscope (Zeiss Axio Zoom V16 or Zeiss Axio Observer Z1) or confocal microscope (Leica TCS SP8). The images were acquired with Zen 2 Pro software and analyzed with Fiji software. Figures were created with Adobe Photoshop and Fiji software.

### Cell culture, shRNA and OSS

Primary hdLECs were cultured on fibronectin-coated six-well plates in EBM-2 MV media and used at passage 6-7 for all experiments. To knock down eNOS, cells were infected with lentiviral particles expressing an shRNA targeting *NOS3* or a control scramble construct expressing GFP for 48 hours. The lentiviral sh*NOS3* (VectorBuilder Inc.) target sequence is CCGGAACAGCACAAGAGTTAT.

The cells were exposed to OSS when they reached 95% confluency. Cells cultured in 6-well plates were exposed to OSS using a test tube rocker (Thermolyne Speci-Mix aliquot mixer model M71015, Barnstead International) with a preset frequency of 18 rpm, 0.3Hz^11^. Cells cultured in 10cm plates were exposed to OSS using a horizontal bidirectional shaker (MS-NOR-30, Major Science) that switched between clockwise and counterclockwise rotation at a frequency of 100rpm, 0.5Hz^19^. All OSS experiments were performed inside a humidified cell culture incubator with 5% CO_2_.

For experiments detecting phospho-eNOS, hdLECs were cultured to reach over 95% confluency after which the medium was switched from EBM-2 MV containing 5% serum to 0.5% serum medium for starvation. After 6 hours of starvation, cells were exposed to OSS for the indicated time points after which lysates were collected. For the SC-79 treatment experiment, cells were starved for 6 hours and then treated with 8μg/mL SC-79 for 30 minutes which was previously reported to lead to increased phospho-AKT levels in hdLECs^13^. DMSO was used as a control.

For L-NAME (100µM and 1mM), L-NMMA (100µM), and KT5823 (5µM and 10µM) inhibitors, hdLECs were cultured to reach over 95% confluency and then the indicated drugs were added to the media. Cells were exposed to static or OSS conditions for 48 hours after which, protein lysates were collected for western blot. For L-NAME and L-NMMA, water was used as a vehicle control. For KT5823, DMSO was used as a vehicle control.

### Western blot

hdLEC lysates were collected using RIPA buffer (Pierce^TM^, Thermo Scientific^TM^) and the total protein concentration was measured using a BCA Protein Assay Kit (Pierce^TM^, Thermo Scientific^TM^). Protein gel electrophoresis was performed with a 4%-12% Blot^TM^ Bis Tris Plus polyacrylamide gel and the Invitrogen^TM^ Mini Gel Tank using 20-25μg of lysate per lane. Dry transfer onto a PVDF membrane was performed using the Invitrogen^TM^ iBlot^TM^ 2 Gel Transfer Device. For blots containing phospho-eNOS, wet transfer onto a PVDF membrane was performed using the Invitrogen^TM^ Mini Blot Module and Invitrogen^TM^ Bolt^TM^ Transfer Buffer. The iBind^TM^ Western Device was used to incubate the membrane with primary and secondary antibodies. The SuperSignal^TM^ West Pico PLUS Chemiluminescent Substrate (Thermo Scientific^TM^) was used to develop the membrane for protein visualization using the Bio-rad ChemiDoc imaging system.

### Co-Immunoprecipitation (Co-IP)

The Co-IP procedure was carried out using the Pierce^TM^ Co-IP kit (Thermo Scientific^TM^). hdLECs lysates were collected using IP/Lysis Wash Buffer (Pierce^TM^, Thermo Scientific^TM^) and protein concentration was measured using a BCA Protein Assay Kit (Pierce^TM^, Thermo Scientific^TM^). For each Co-IP reaction, an antibody-coupled resin was prepared in a Pierce^TM^ Spin Column by mixing 50ug of antibody with the AminoLink^TM^ Plus Coupling Resin and incubating on a rotator (Labnet^TM^ Revolver^TM^ Tube Mixer) at room temperature for 2 hours. Lysate (1mg) was pre-cleared by incubating with the Pierce^TM^ Control Agarose Resin at 4µC for 30 minutes with constant rotation on a rotator. The supernatant was then incubated with the antibody-coupled resin at 4µC overnight with constant rotation on a rotator. The next day, the samples were washed 4 times with IP/Lysis Wash Buffer (Pierce^TM^, Thermo Scientific^TM^). The protein complex was then eluted using the elution buffer provided in the kit and western blot with dry protein transfer was performed as described above.

### Statistics

All data are represented as mean ± SEM. An unpaired 2-sided Student’s *t-*test was performed to determine significant differences between 2 groups. p<0.05 was considered significant. Graphpad Prism software (version 6) was used for all statistical analyses and to plot quantitative data.

## RESULTS

### eNOS is upregulated and activated in lymphatic valve LECs

To investigate eNOS expression in developing and mature valves, we harvested mesenteries from WT embryos and performed whole-mount immunostaining for PROX1 and eNOS. At E16.5, many PROX1^high^ valve specification clusters were observed in the mesentery as previously reported^9^ and eNOS was highly expressed in these PROX1^high^ valve LECs compared to the surrounding PROX1^low^ lymphangion LECs (Supplemental Figure 1, A-F). At later developmental stages, eNOS was highly expressed in the LECs of mature valves at E18.5 (Figure 1, A-F), P3 (Supplemental Figure 1, G-L) and P7 (Supplemental Figure 1, M-R).

**Fig 1:**
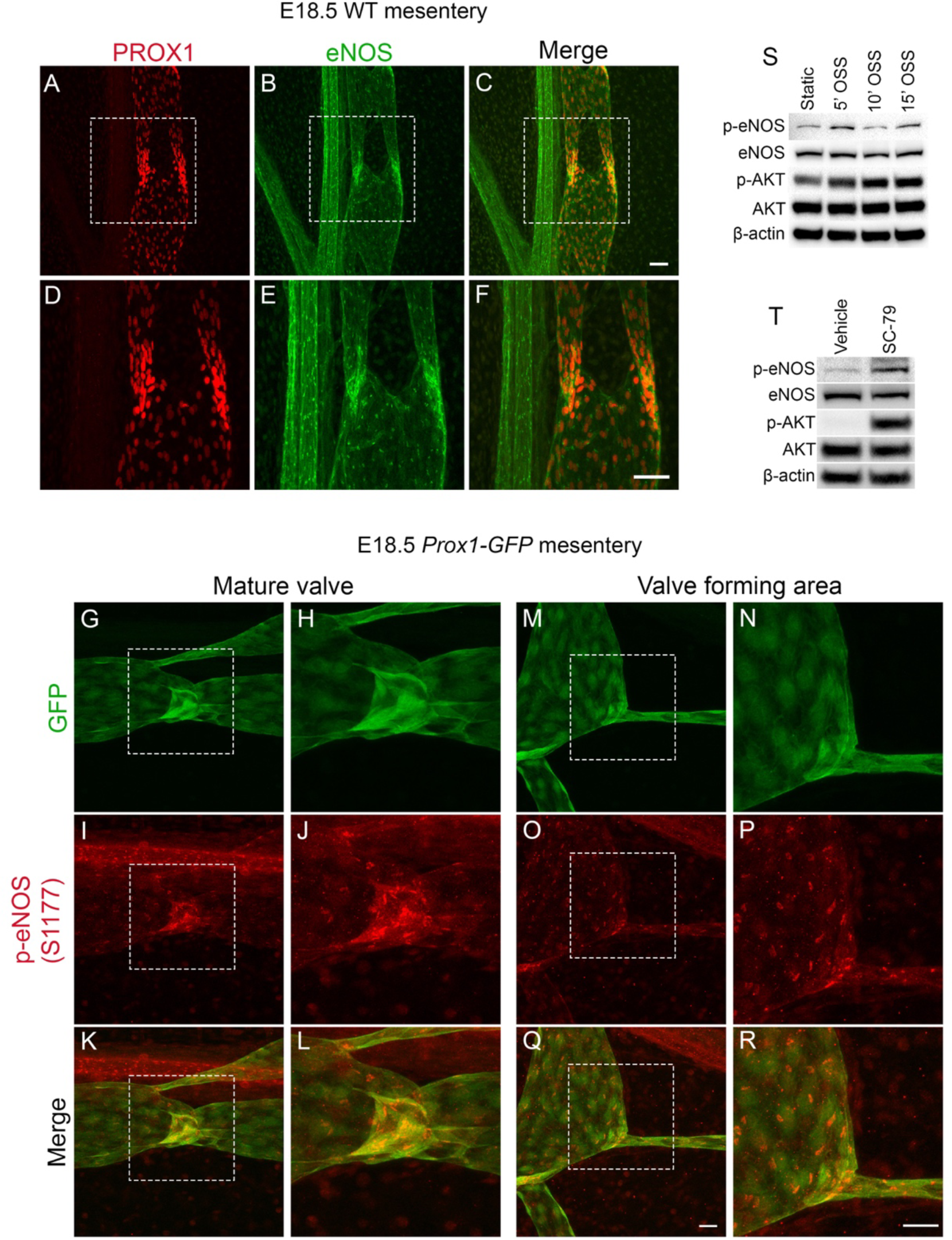
eNOS is upregulated and activated in response to OSS. (A-F) Whole-mount immunostaining of PROX1 (red) and eNOS (green) in WT mesenteries at E18.5. D, E and F are high-magnification images of the white boxed insets in A, B and C respectively. (G-R) Whole-mount immunostaining of GFP (green) and phospho-Ser1177 eNOS (red) in WT mesenteries at E18.5 showing a (G-L) mature valve and (M-R) valve forming area. H, J, L, N, P and R are high-magnification images of the white boxed insets in G, I, K, M, O and Q respectively. (S) Western blot for indicated proteins using lysates from hdLECS exposed to static conditions or OSS conditions for 5, 10 and 15 mins. (T) Western blot for indicated proteins using lysates from hdLECS treated with vehicle or SC-79. Scale bars are 50µm in A-F and 25µm in G-R.

Although eNOS is highly expressed in developing and mature valve LECs, it is unknown whether eNOS is phosphorylated and activated in response to OSS during embryonic valve specification. To address this, we analyzed mesenteries at E18.5, a timepoint when fully mature valves can be found, from *Prox1-GFP* reporter mice that allow convenient visualization of lymphatic valve morphology^15^. Whole-mount immunostaining for phospho-Ser1177 of eNOS revealed strong phospho-eNOS staining in the GFP^high^ LECs of mature valves (Figure 1, G-L) and valve forming areas (Figure 1, M-R). To investigate whether OSS induces phosphorylation of eNOS in LECs, we exposed cultured human dermal LEC (hdLECs) to OSS for 5, 10 and 15 minutes or static conditions as a control and performed western blot. We found that eNOS phosphorylation peaked at 5 and 15 minutes of OSS, whereas AKT phosphorylation steadily increased from 5 to 15 minutes of OSS (Figure 1S). To confirm that eNOS is a downstream target of AKT, we treated hdLECs with the AKT activator SC-79 or vehicle and performed western blot. We found that SC-79 treatment led to an increase in phosphorylated AKT and phosphorylated eNOS (Figure 1T). These results suggest that eNOS is activated by AKT in response to OSS.

### eNOS regulates lymphatic valve development during embryogenesis

To investigate whether eNOS regulates valve specification, we performed whole-mount immunostaining for PROX1 and FOXC2 in WT and *Nos3^−/−^* mesenteries at E16.5. We observed several cell clusters expressing high levels of PROX1 and FOXC2 in WT mesenteries (Figure 2, A-C) and these were reduced in *Nos3^−/−^* mesenteries (Figure 2, D-F). Since FOXC2 expression is high in the adjacent arterial smooth muscles^20^, we used PROX1 staining to quantify specification clusters in both groups and normalized the number to vessel length. We found 45% fewer PROX1^high^ clusters per millimeter in *Nos3^−/−^* mesenteries compared to WT controls (Figure 2G) while there was no significant difference in the total length of lymphatic vessels between the two groups (Figure 2H).

**Fig 2:**
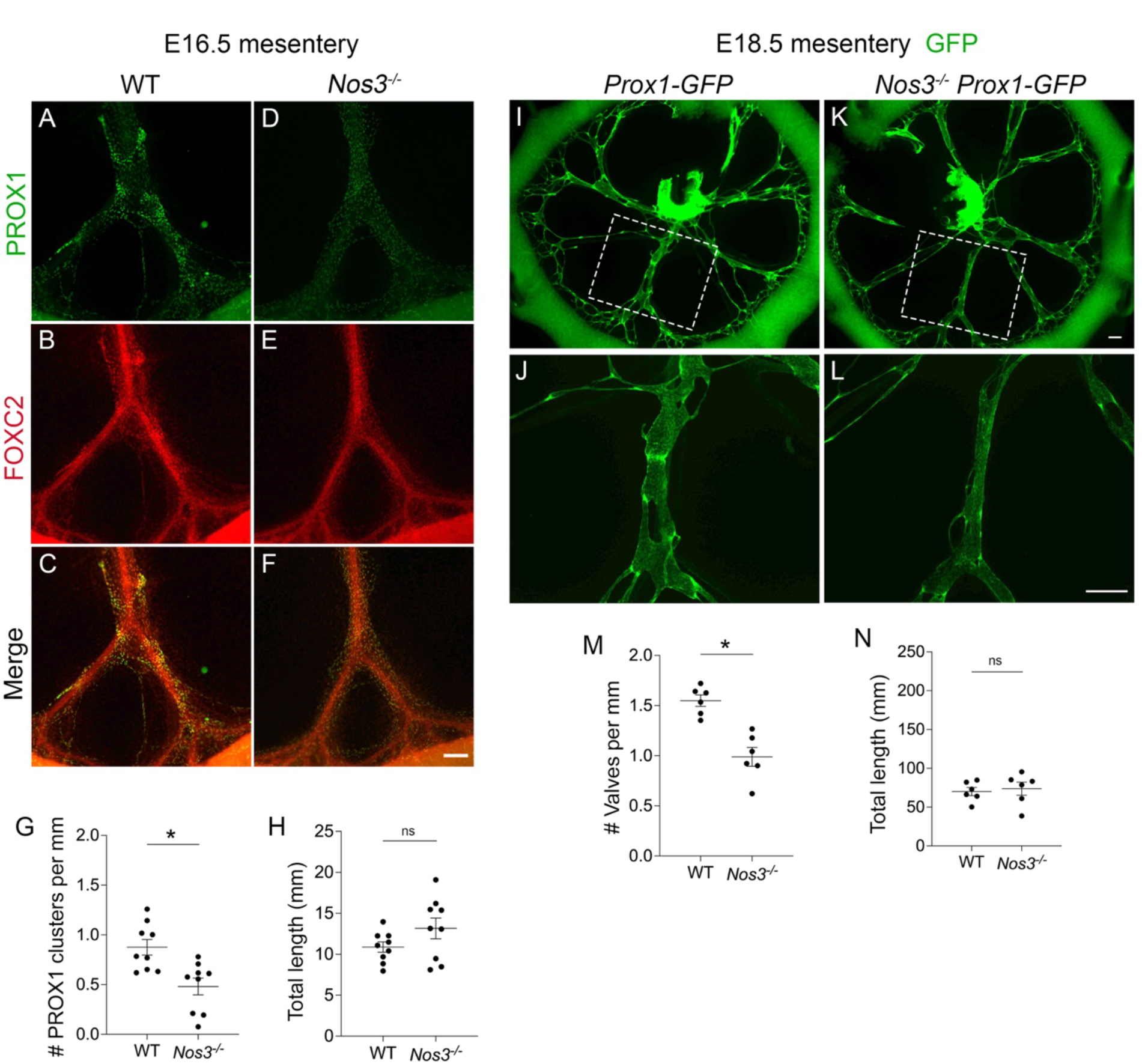
*Nos3^−/−^* embryos have a defect in lymphatic valve development. (A-F) Whole-mount immunostaining of PROX1 (green) and FOXC2 (red) in WT and *Nos3^−/−^* E16.5 mesenteries. (G) PROX1 clusters per millimeter from each E16.5 mesentery. (H) Total length of lymphatic vessels from each E16.5 mesentery. (G,H) All values are means ± SEM of n=9 littermates per genotype. (I-L) Fluorescence imaging of Prox1-GFP (green) mesenteries at E18.5. (J,L) High-magnification images of the white boxed areas in I and K respectively. (M) Valves per millimeter from each E18.5 mesentery. (N) Total length of lymphatic vessels from each E18.5 mesentery. (M,N) All values are means ± SEM of n=6 littermates per genotype. *P<0.05, unpaired Student’s *t*-test. Scale bars are 150µm in A-F and 300µm in I-L.

Next, we wanted to determine whether the valve specification defects led to fewer mature lymphatic valves at E18.5, a timepoint when fully formed valves first appear in the mesentery. We harvested mesenteries from *Nos3^−/−^;Prox1-GFP* embryos and *Prox1-GFP* controls at E18.5 and counted the GFP^high^ lymphatic valves in both groups (Figure 2, I-L). Quantification of the GFP^high^ lymphatic valves revealed that *Nos3^−/−^;Prox1-GFP* mice had 36% fewer valves per millimeter compared to controls (Figure 2M) and there was no significant difference in the total length of lymphatic vessels between the two groups (Figure 2N).

### *Nos3^−/−^* mice have fewer lymphatic valves at P3

To investigate whether the valve specification defects lead to fewer valves postnatally, we quantified all of the GFP^high^ valves in *Nos3^−/−^;Prox1-GFP* and *Prox1-GFP* control mesenteries at postnatal day P3. As expected, we found that *Nos3^−/−^* mice had 60% fewer total valves compared to controls (Figure 3, A-D and I) while there was no significant difference in the total length of lymphatic vessels between the two groups (Figure 3J).

**Fig 3:**
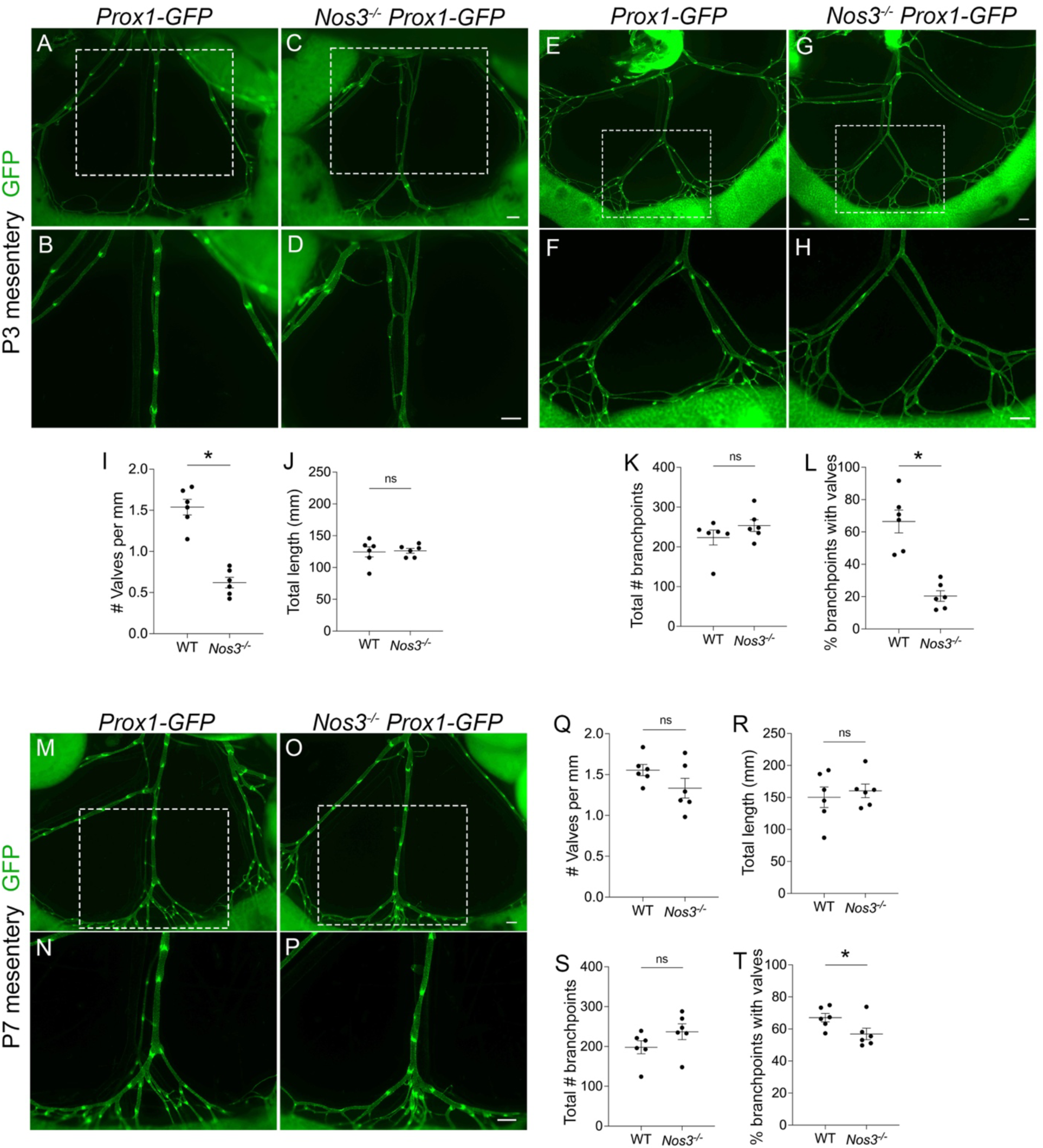
*Nos3^−/−^* animals have fewer lymphatic valves postnatally. (A-H) Fluorescence imaging of Prox1-GFP (green) mesenteries at P3. B, D, F and H are high-magnification images of the white boxed insets in A, C, E and G. (I) Valves per millimeter from each mesentery at P3. (J) Total length of lymphatic vessels from each mesentery at P3. (K) Total number of branchpoints from each mesentery at P3. (L) Percentage of total branchpoints containing valves from each mesentery at P3. (M-P) Fluorescence imaging of Prox1-GFP (green) mesenteries at P7. (N,P) High-magnification images of the white boxed insets in M and O respectively. (Q) Valves per millimeter from each mesentery at P7. (R) Total length of lymphatic vessels from each mesentery at P7. (S) Total number of branchpoints from each mesentery at P7. (T) Percentage of total branchpoints containing valves from each mesentery at P7. All values are means ± SEM of n=6 littermates per genotype. *P<0.05, unpaired Student’s *t*-test. Scale bars are 300µm in A-H and M-P.

Since lymphatic valves preferentially grow in areas of lymph flow reversal such as branchpoints^9^, we quantified the total branchpoints and the branchpoints containing valves in *Nos3^−/−^;Prox1-GFP* mice and *Prox1-GFP* controls. We found that there was no significant difference in the total number of branchpoints between the two groups (Figure 3, E-H and K), indicating that *Nos3^−/−^* mice did not have a defect in vessel branching. In contrast, only 20% of the branchpoints in *Nos3^−/−^;Prox1-GFP* mice had valves, which was significantly less than the 66% of branchpoints that had valves in controls (Figure 3L).

To determine whether the valve defects persisted at even later timepoints, we harvested mesenteries from *Nos3^−/−^;Prox1-GFP* mice and *Prox1-GFP* controls at P7 and quantified valves. Unexpectedly, we found that there was no significant difference in valve number and total length of lymphatic vessels between the two groups (Figure 3, M-R). At P7, 57% of branchpoints in *Nos3^−/−^;Prox1-GFP* mice had valves and this was significantly lower than controls, in which 67% of branchpoints had valves (Figure 3T), and there was no significant difference in the total number of branchpoints between the two groups (Figure 3S). Interestingly, the percentage of branchpoints with valves in *Nos3^−/−^;Prox1-GFP* mice increased from 20% at P3 to 57% at P7, suggesting that *Nos3^−/−^* mice experience a delay in valve development.

### eNOS regulates expression of the valve genes *Prox1* and *Foxc2*

Valve LECs are specified when OSS activates signaling pathways that lead to upregulation of PROX1 and FOXC2^9, 11^. Since *Nos3^−/−^* mice experience a defect in valve specification, we wanted to investigate whether eNOS regulated the expression of PROX1 and FOXC2. We harvested mesenteries from *Nos3^−/−^* mice and WT littermate controls at P3, performed whole-mount immunostaining for PROX1, FOXC2 and VE-cadherin and took high magnification confocal images of the lymphangion and valve areas. We observed that *Nos3^−/−^* mice had reduced PROX1 and FOXC2 expression compared to WT controls in the lymphangion LECs (Figure 4, A-F). Interestingly, we observed no change in expression of PROX1 and FOXC2 in the valve LECs between the two groups (Figure 4, G-L). When the pixel intensity was quantified, it revealed a significant decrease in PROX1 and FOXC2 expression in the lymphangion LECs in *Nos3^−/−^* mice compared to WT controls (Figure 4, M and N). Conversely, there was no significant difference in PROX1 and FOXC2 expression in valve LECs between the two groups (Figure 4, O and P).

**Fig 4:**
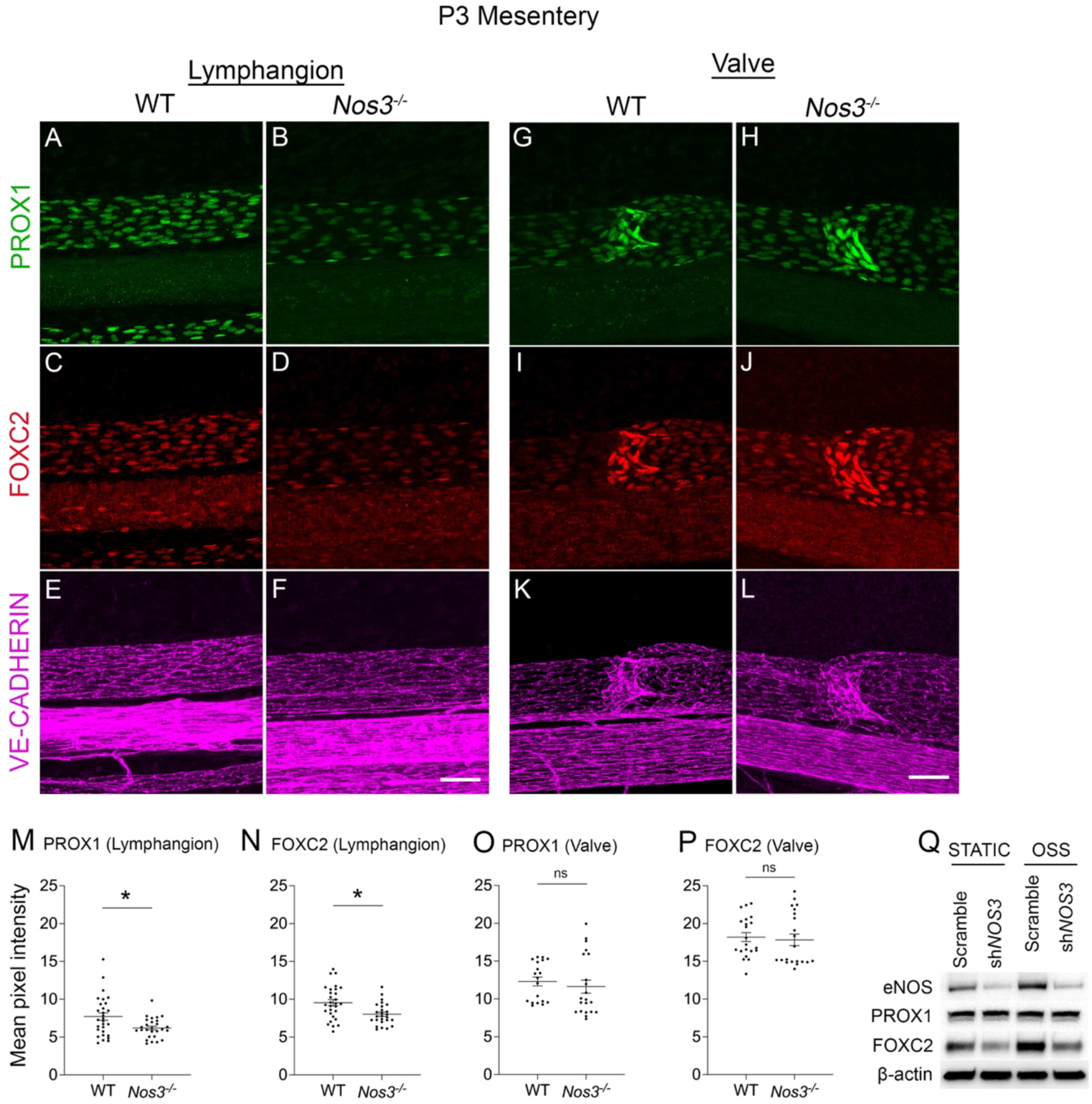
eNOS regulates PROX1 and FOXC2 expression. (A-L) Whole-mount immunostaining of PROX1 (green), FOXC2 (red) and VE-Cadherin (magenta) in WT and *Nos3^−/−^* P3 mesenteries. (M,N) Mean pixel intensity measurements of (M) PROX1 and (N) FOXC2 in lymphangion LECs from each mesentery. (O,P) Mean pixel intensity measurements of (O) PROX1 and (P) FOXC2 in valve LECs from each mesentery. (Q) Western blot for indicated proteins using lysates from hdLECS cultured under static or OSS conditions for 48 hours and transfected with a control scramble or shRNA against *NOS3.* All values are means ± SEM of n=4-6 lymphangion or valve areas from N=5-6 littermates per genotype. *P<0.05, unpaired Student’s *t*-test. Scale bars are 50µm in A-L.

Next, we infected cultured hdLECs with a lentivirus expressing an shRNA against *NOS3* or a scramble control shRNA and exposed them to static or OSS conditions. Western blot analysis confirmed efficient knockdown of eNOS protein in the sh*NOS3* LECs and showed that eNOS and FOXC2 expression was increased in scramble LECs in response to OSS as previously reported^9^. In support of our *in vivo* data, sh*NOS3* LECs had reduced FOXC2 expression compared to scramble-infected LECs in both static and OSS conditions. PROX1 expression did not change in response to OSS in scramble-infected LECs as previously published^11^ and was not affected by *NOS3* knockdown (Figure 4Q).

### eNOS regulates β-catenin signaling

Valve specification is regulated by the transcription factor β-catenin, which directly binds the promoter regions of *Prox1* and *Foxc2* and upregulates their expression in response to OSS^11^. Since our data showed that eNOS regulates PROX1 and FOXC2 expression, we investigated whether eNOS regulates β-catenin signaling. In endothelial cells, β-catenin signaling is regulated by a multiprotein complex which phosphorylates cytoplasmic β-catenin and consequently marks it for proteasomal degradation, thereby limiting the fraction of β-catenin that can translocate to the nucleus and transcribe target genes^21^. We harvested mesenteries from *Nos3^−/−^* and WT embryos at E18.5 and performed whole-mount immunostaining using antibodies against PROX1 and non-phosphorylated β-catenin, which represents the pool of β-catenin that is spared by the destruction complex and is free to transcribe target genes. We found that in WT embryos, non-phosphorylated β-catenin localized at the cell membrane and colocalized with PROX1 in the nucleus (Figure 5, A-F). Conversely, in *Nos3^−/−^* embryos, non-phosphorylated β-catenin was mainly localized at the cell membrane with no visible nuclear staining (Figure 5, G-L). Immunostaining of hdLECs showed that under static conditions there was no detectable β-catenin in sh*NOS3* hdLECs compared to scramble hdLECs (Figure 5 M and O). Scramble hdLECs exposed to OSS led to increased accumulation of nuclear β-catenin compared to static conditions as previously reported (Figure 5N)^11^. In support of our *in vivo* data, sh*NOS3* hdLECs exposed to OSS showed no visible nuclear β-catenin staining (Figure 5P), indicating that eNOS is responsible for the nuclear translocation of β-catenin.

**Fig 5:**
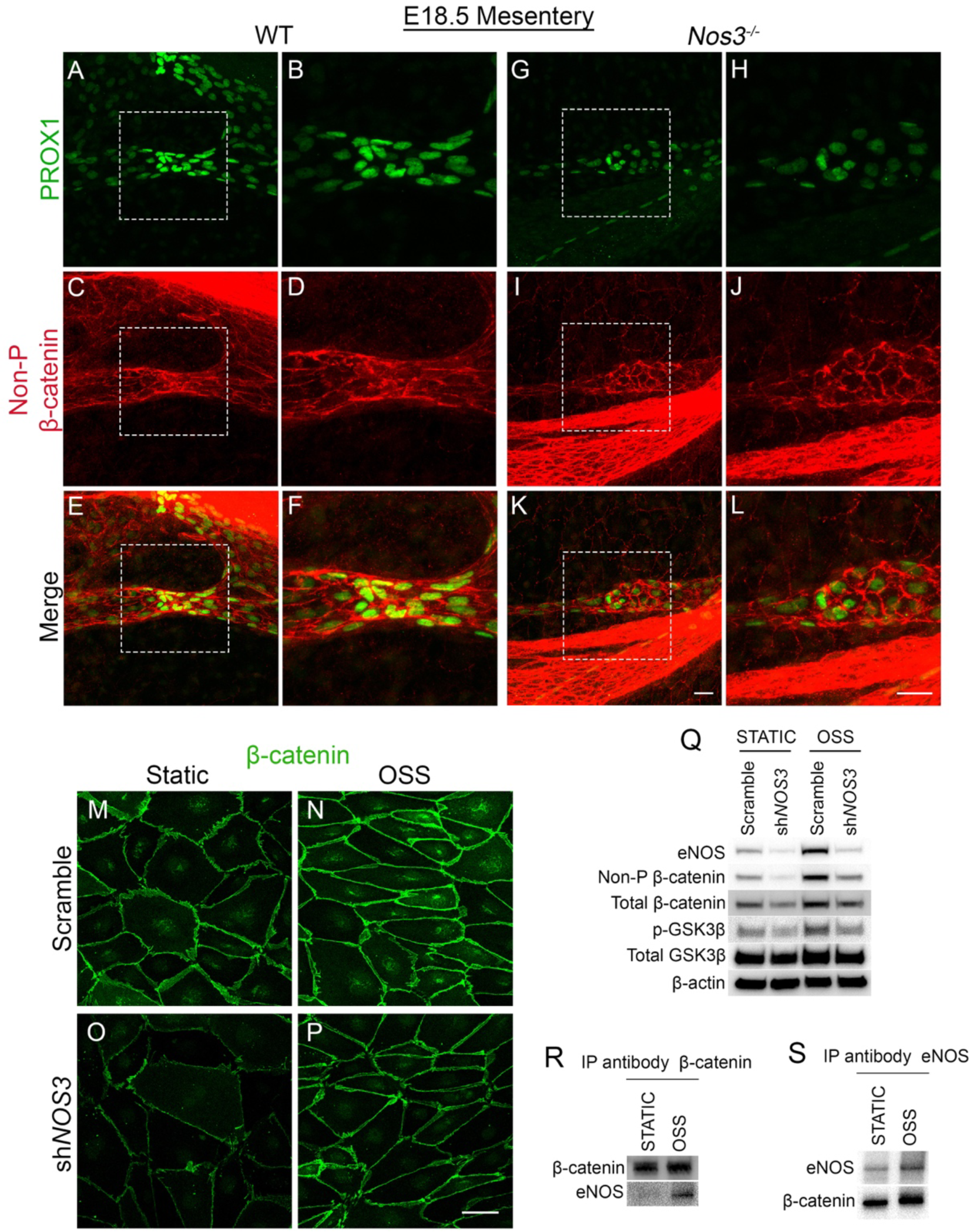
eNOS regulates β-catenin signaling. (A-L) Whole-mount immunostaining of PROX1 (green) and Non-phosphorylated β-catenin (referred to as Non-P β-catenin) (red) in WT and *Nos3^−/−^* E18.5 mesenteries. B, D, F, H, J and L are high-magnification images of the white boxed areas in A, C, E, G, I and K respectively. (M-P) Immunostaining of β-catenin (green) in hdLECs cultured under static or OSS conditions for 48 hours and transfected with a control scramble or shRNA against *NOS3.* (Q) Western blot for indicated proteins using lysates from scramble or sh*NOS3* hdLECS cultured under static or OSS conditions for 48 hours. (R,S) Lysates from hdLECS cultured under static or OSS conditions were immunoprecipitated (IP) using (R) anti-β-catenin antibody or (S) anti-eNOS antibody followed by western blot for indicated proteins. Scale bars are 25µm in A-L.

Because phosphorylation and subsequent degradation of β-catenin inhibits its transcriptional activity^21^, we performed western blot for non-phosphorylated β-catenin and total β-catenin on lysates collected from scramble and sh*NOS3* hdLECs exposed to static or OSS. In response to OSS, scramble LECs showed increased non-phosphorylated β-catenin and total β-catenin levels as previously published^11^. In support of our *in vivo* data, we found that *NOS3*-knockdown LECs showed decreased non-phosphorylated β-catenin and total β-catenin levels compared to scramble LECs, in both static and OSS conditions (Figure 5Q), indicating that in the absence of eNOS, there is more degradation of β-catenin that blunts the response to flow.

### eNOS regulates GSK3β activity

Next, we investigated the mechanism by which eNOS regulates β-catenin signaling. A study in HUVECs showed that eNOS activation and subsequent nitric oxide (NO) production can enhance nuclear translocation of β-catenin^22^. GSK3β is part of the β-catenin destruction complex, and the study showed that NO-mediated activation of protein kinase G (PKG) resulted in phosphorylation of GSK3β, which inhibits GSK3β activity^21, 22^. To test whether eNOS regulates GSK3β activity in hdLECs, we performed western blot for phospho-GSK3β. In scramble hdLECs, we found that GSK3β is phosphorylated and thus inhibited in response to OSS. Furthermore, we found that *NOS3*-knockdown hdLECs showed reduced phospho-GSK3β levels compared to scramble hdLECs in both static and OSS conditions (Figure 5Q). This suggests that GSK3β could be degrading β-catenin in the absence of eNOS.

### eNOS regulates β-catenin signaling independent of NO production

Since we observed phospho-eNOS in valve LECs, we tested whether eNOS regulated β-catenin signaling via NO production. We exposed hdLECs to static or OSS conditions in the presence of L-N^G^-Nitro arginine methyl ester (L-NAME), a pan-NOS inhibitor, or vehicle control. Western blot analysis showed no change in levels of non-phosphorylated β-catenin, total β-catenin, phospho-GSK3β and total GSK3β in hdLECs treated with 100µM and 1mM L-NAME compared to vehicle-treated hdLECs, in both static and OSS conditions (Supplemental Figure 2 A and B). We also treated hdLECs with an irreversible pan-NOS inhibitor, L-N^G^-Monomethyl Arginine (L-NMMA) and consistently found no change in non-phosphorylated β-catenin, total β-catenin, phospho-GSK3β and total GSK3β between vehicle and 100µM LNMMA-treated hdLECs, in both static and OSS conditions (Supplemental Figure 2C).

To test whether eNOS regulates β-catenin signaling through NO-PKG signaling, we treated hdLECs with a PKG inhibitor, KT5823 or vehicle and exposed them to static or OSS conditions. Western blot analysis showed that in static conditions there was no change in total β-catenin, non-phosphorylated β-catenin, phospho-GSK3β and total GSK3β levels between 5µM or 10µM KT5823-treated hdLECs and vehicle-treated hdLECs (Supplemental Figure 2D). In OSS conditions, there was no change in total β-catenin, phospho-GSK3β and total GSK3β levels between 5µM or 10µM KT5823-treated hdLECs and vehicle-treated hdLECs (Supplemental Figure 2D). Non-phosphorylated β-catenin levels did not change in 5µM KT5823-treated hdLECs but slightly increased in 10µM KT5823-treated hdLECs compared to vehicle-treated hdLECs (Supplemental Figure 2D). This is likely an off-target effect of the drug since the 10µM dosage is approximately 200 times higher than the IC_50_.

### eNOS forms a complex with β-catenin in hdLECs

Since loss of eNOS protein but not NO inhibition was found to affect β-catenin, we decided to test whether eNOS forms a complex with β-catenin to regulate its signaling. Evidence for eNOS binding β-catenin has been reported in HUVECs but has not been investigated in LECs^22^. We immunoprecipitated β-catenin from hdLEC lysates that were exposed to static or OSS conditions (Figure 5R). A co-IP western blot revealed the presence of β-catenin under static and OSS conditions as a positive control. While eNOS protein was barely present in the static condition, its presence was enhanced by OSS, suggesting more binding of these two proteins in flow. We performed the reverse Co-IP blot using an eNOS antibody as a probe (Figure 5S), which revealed that β-catenin binds eNOS under static conditions and the binding is enhanced by OSS. These data suggest that eNOS forms a complex with β-catenin to regulate its activity.

### Postnatal deletion of β-catenin does not lead to a complete loss of lymphatic valves

Embryonic lymphatic-specific deletion of β-catenin has been previously shown to result in a complete loss of valve specification clusters at E16.5 and mature valves at E18.5^11^. However, the role of β-catenin in postnatal lymphatic valve development has not been investigated. To delete β-catenin in a lymphatic-specific manner postnatally, we generated a transgenic mouse line expressing tamoxifen-inducible Cre recombinase (*CreER^T2^)* that is under the control of the *Prox1* gene promoter. This strain was interbred with *Ctnnb1^flox/flox^* and *Prox1-GFP* mice to generate *Prox1CreER^T2^;Ctnnb1^flox/flox^;Prox1-GFP* mice (hereafter: *Ctnnb1^LEC-KO^* (JScal)) and control littermates lacking Cre recombinase (*Ctnnb1^fl/fl^)*^16^. We administered tamoxifen (TM) at P1 and P3 and analyzed the mesentery at P8 (Figure 6A). Whole-mount immunostaining for β-catenin revealed ∼100% efficient deletion of the protein from lymphatic vessels (Figure 6B-E). We further verified efficiency and specificity of the *Prox1CreER^T2^* (JScal) line by crossing it with the *R26-mTmG* reporter and analyzing reporter gene expression following TM administration. Cre-mediated GFP expression was restricted to the lymphatic vasculature and other Prox1-expressing tissues (data not shown). We found that *Ctnnb1^LEC-KO^* (JScal) mice had 58% fewer lymphatic valves at P8 compared to controls (Figure 6 F-I and N) while there was no significant difference in the total length of lymphatic vessels between the two groups (Figure 6Q). To further validate our *Prox1CreER^T2^* strain, we crossed the *Ctnnb1^flox/flox^* and *Prox1-GFP* mice to a previously published *Prox1CreER^T2^* strain^17^ to again generate *Prox1CreER^T2^;Ctnnb1^flox/flox^;Prox1-GFP* mice (hereafter: *Ctnnb1^LEC-KO^* (TMak)) and control littermates lacking Cre recombinase (*Ctnnb1^fl/fl^*). Following TM administration at P1 and P3, we analyzed the mesentery at P8 and found that *Ctnnb1^LEC-KO^* (TMak) mice had 55% fewer lymphatic valves compared to controls (Figure 6J-M and O) while there was no significant difference in the total length of lymphatic vessels between the two groups (Figure 6R). There was no significant difference in the valve number and length of lymphatic vessels between *Ctnnb1^LEC-KO^* (JScal) and *Ctnnb1^LEC-KO^* (TMak) mice (Figure 6P and S). In contrast to the requirement of β-catenin in embryonic specification, our data show that β-catenin maintains approximately 50% of postnatal valves.

**Fig 6:**
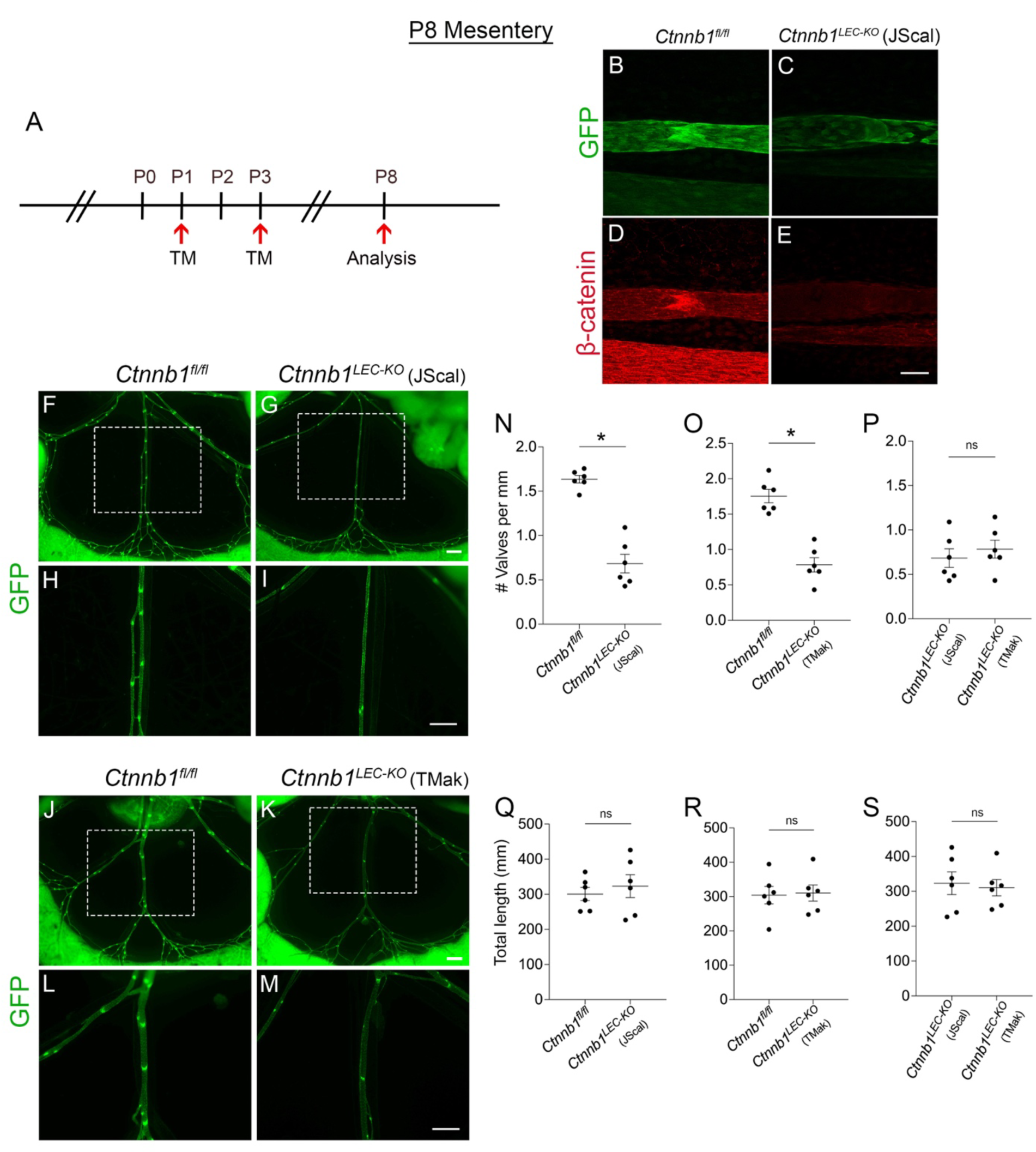
Postnatal β-catenin deletion does not lead to a complete loss of valves. (A) Tamoxifen injection schedule for postnatal deletion of *Ctnnb1.* (B-E) Whole-mount immunostaining of GFP (green) and β-catenin (red) in *Ctnnb1^fl/fl^* and *Ctnnb1^LEC-KO^* (JScal) P8 mesenteries. (F-M) Fluorescence imaging of Prox1-GFP (green) mesenteries at P8. H, I, L and M are high-magnification images of the white boxed areas in F, G, J and K respectively. (N-P) Valves per millimeter from each mesentery at P8. (Q-S) Total length of lymphatic vessels from each mesentery at P8. All values are means ± SEM of n=6 littermates per genotype. *P<0.05, unpaired Student’s *t*-test. Scale bars are 50µm in B-E and 500µm F-M.

### β-catenin gain-of-function allele does not rescue valve loss in *Nos3^−/−^* mice

Since our data suggests that eNOS regulates β-catenin signaling to regulate lymphatic valve development, we decided to test whether enhancing β-catenin signaling could rescue the valve defects in *Nos3^−/−^* mice. To genetically enhance β-catenin signaling, we utilized a previously published mouse model in which exon 3 of the β-catenin gene is floxed (*Ctnnb1^lox(ex3)^*)^18^. Deletion of exon 3 prevents β-catenin phosphorylation and targeting for proteasomal degradation, thereby leading to its accumulation and gain-of-function (GOF)^18^. To induce the β-catenin-GOF allele in *Nos3^−/−^* mice in a lymphatic specific manner, we generated *Prox1CreER^T2^;Ctnnb1^+/lox(ex3)^;Nos3^−/−^;Prox1-GFP* mice (hereafter referred to as *Nos3^−/−^;Ctnnb1^LEC-GOF^*). Littermates lacking Cre recombinase (hereafter referred to as *Nos3^−/−^;Ctnnb1^+/lox(ex3)^*) served as controls expected to phenocopy *Nos3^−/−^* mice. First, we induced embryonic expression of the β-catenin-GOF allele at E14.5 and analyzed both groups at E18.5 (Figure 7A). Unexpectedly, we found that β-catenin GOF did not lead to an increase in the number of GFP^high^ valve areas in *Nos3^−/−^* embryos (Figure 7B-E). Quantification revealed no significant difference in valve number between the two groups (Figure 7F) and a 40% increase in length of lymphatic vessels in *Nos3^−/−^; Ctnnb1^LEC-GOF^* compared to *Nos3^−/−^;Ctnnb1^+/lox(ex3)^* mice (Figure 7G). Next, we induced postnatal expression of the β-catenin-GOF allele at P0 and analyzed mesenteries from both groups at P3 (Supplemental Figure 3A). In contrast to our embryonic data, we found that postnatal β-catenin GOF led to a significant loss of valves in *Nos3^−/−^* mice (Supplemental Figure 3 B-E). Quantification revealed a 60% decrease in valve number in *Nos3^−/−^;Ctnnb1^LEC-GOF^* mice compared to *Nos3^−/−^; Ctnnb1^+/lox(ex3)^* mice (Supplemental Figure 3F) with no difference in total length of lymphatic vessels between the two groups (Supplemental Figure 3G).

**Fig 7:**
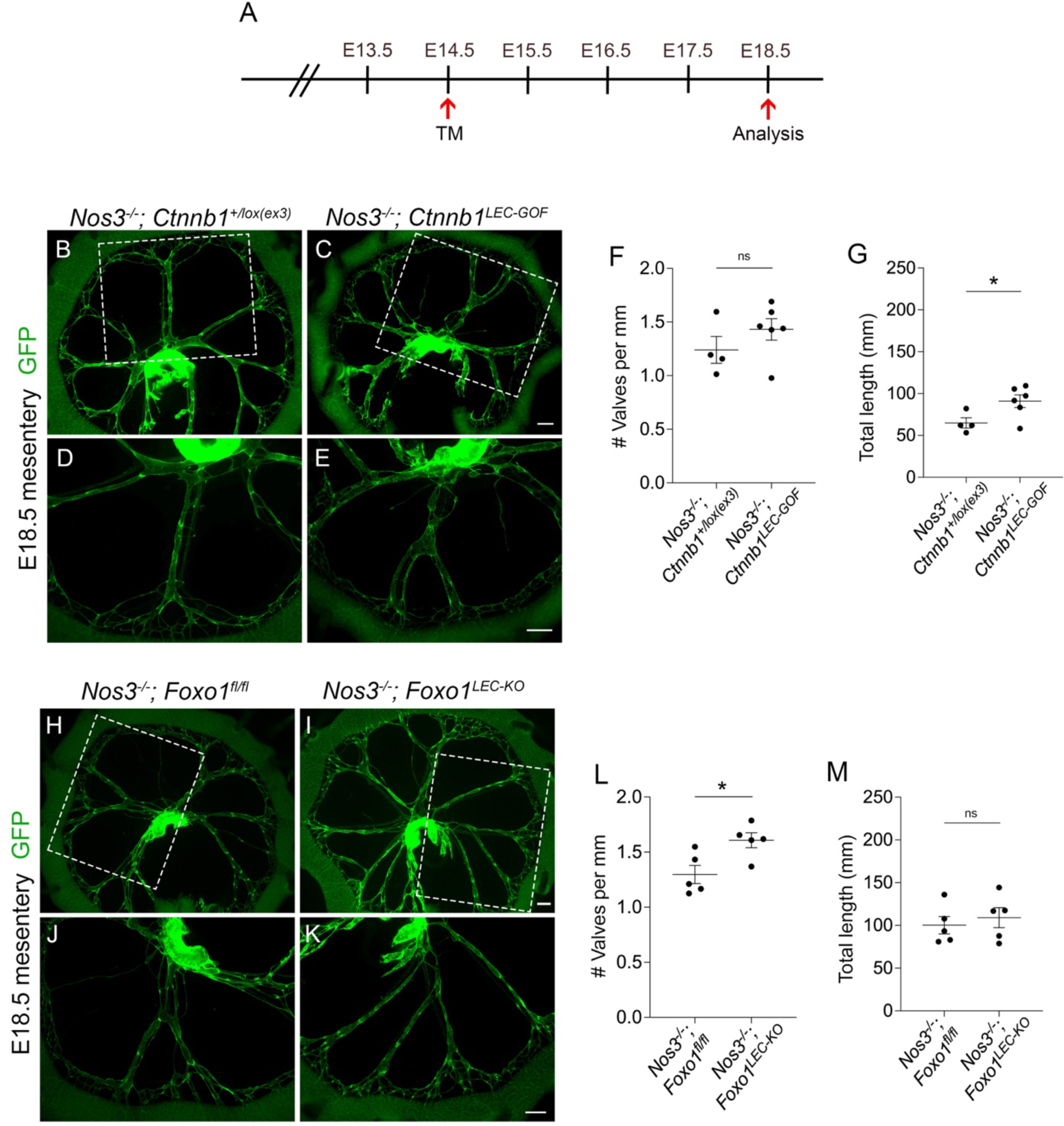
*Foxo1* deletion but not β-catenin GOF rescues valve loss in *Nos3^−/−^* embryos. (A) Tamoxifen injection schedule for embryonic deletion of *Ctnnb1* exon 3 or *Foxo1.* (B-E and H-K) Fluorescence imaging of E18.5 Prox1-GFP (green) mesenteries from (B-E) *Nos3^−/−^;Ctnnb1^LEC-GOF^* and control littermates lacking *Prox1CreER^T2^* and (H-K) *Nos3^−/−^;Foxo1^LEC-KO^* and control littermates lacking *Prox1CreER^T2^*. (F,L) Valves per millimeter from each mesentery at E18.5. (G,M) Total length of lymphatic vessels from each mesentery at E18.5. All values are means ± SEM of n=4-6 littermates per genotype. *P<0.05, unpaired Student’s *t*-test. Scale bars are 500µm in B-E and H-K.

### Foxo1 deletion rescues valve loss in *Nos3^−/−^* embryos

Since our data showed that eNOS regulates the expression of PROX1 and FOXC2, we decided to test whether directly inducing FOXC2 expression could rescue the valve defects in *Nos3^−/−^* mice. FOXO1 is a transcription factor that was shown to repress the expression of valve genes and directly bound the *Foxc2* promoter as a repressor^13^. The same study showed that lymphatic-specific deletion of *Foxo1* stimulated lymphatic valve formation and rescued the loss of valves in *Foxc2* heterozygous mice^13^. Additionally, a recent study showed that inhibition of FOXO1 using the drug AS1842865 increased the nuclear translocation of β-catenin in hdLECs through an unidentified mechanism^23^. Thus, we generated *Prox1CreER^T2^;Foxo1^fl/fl^; Nos3^−/−^;Prox1-GFP* mice (hereafter: *Nos3^−/−^;Foxo1^LEC-KO^*). Littermates lacking Cre recombinase (hereafter: *Nos3^−/−^; Foxo1^fl/fl^*) served as controls expected to phenocopy *Nos3^−/−^* mice. Embryonic deletion of *Foxo1* (Figure 7A) led to a significant increase in GFP^high^ valve areas in *Nos3^−/−^* embryos (Figure 7H-K). Quantification revealed an increase in valve number in *Nos3^−/−^;Foxo1^LEC-KO^* mice back to WT control levels in Figure 2M (∼ 1.5 valves per mm) (Figure 7L). There was no difference in the total length of lymphatic vessels between *Nos3^−/−^;Foxo1^LEC-KO^* and *Nos3^−/−^;Foxo1^fl/fl^* (Figure 7M).

## DISCUSSION

OSS generated by lymph flow activates mechanotransduction signaling pathways to regulate lymphatic valve development^9–13^. However, the exact signaling pathways that are activated by OSS have not been fully identified. As eNOS is a well-studied shear stress signaling molecule in BECs, we investigated its role in lymphatic valve development. Previous work showed that eNOS expression is induced in LECs shortly after the onset of lymph flow^9^. Consistent with this study, we show that eNOS is upregulated in the LECs of valve specification clusters and continues to be highly expressed in developing and mature valves. Furthermore, we show that eNOS regulates lymphatic valve specification by forming a complex with the transcription factor β-catenin to regulate its nuclear translocation and binding to the *Prox1* and *Foxc2* promoters.

### eNOS regulates expression of the valve genes PROX1 and FOXC2

LEC clusters that are specified to form lymphatic valves express high levels of PROX1 and FOXC2 compared to the surrounding LECs and are first observed at E16.5 in the mouse mesentery^9^. We found that *Nos3^−/−^* mice experience a defect in valve specification that led to fewer valves at E18.5 and P3. Since lymphatic valves preferentially grow at branchpoints, we confirmed that *Nos3^−/−^* mice have a valve development defect and not a branching defect by showing that *Nos3^−/−^* mice had normal branching but failed to grow valves at the branchpoints.

Consistent with a defect in valve specification, we found that *Nos3^−/−^* mice have significantly reduced levels of PROX1 and FOXC2 in the lymphangion LECs compared to WT controls. This was recapitulated *in vitro* where *NOS3* knockdown led to reduced FOXC2 expression at the protein level. *NOS3* knockdown did not affect PROX1 levels *in vitro,* however this was not surprising because previous reports have established that the *in vitro* OSS model does not modulate PROX1 expression^11–13^. Altogether, our data indicate that *Nos3^−/−^* LECs have a blunted ability to upregulate PROX1 and FOXC2 in response to OSS to specify valve LECs. Interestingly, we saw that there was no difference in PROX1 and FOXC2 expression levels in the LECs of fully formed valves in WT and *Nos3^−/−^* mice. We reason that LECs of mature valves must have adequate levels of PROX1 and FOXC2 if they were able to form a valve. Other signaling pathways that can modulate PROX1 and FOXC2 expression independently of eNOS that may give rise to the limited number of valves that we observed in the *Nos3^−/−^* mice^13^.

### eNOS regulates lymphatic valve development independent of NO production

In BECs, AKT has been shown to phosphorylate the Serine1177 residue of eNOS in response to fluid shear stress, which leads to its activation and subsequent NO production^14^. Moreover, we previously published that OSS induced AKT activation in cultured hdLECs and injecting an AKT activator in WT mice led to ectopic lymphatic valve growth in the mice^12^. Based on this we initially developed the hypothesis that eNOS-NO signaling acts downstream of AKT activation in regulating lymphatic valve development. We detected phospho-Ser1177 eNOS in the LECs of developing and mature valves *in vivo* and found that in cultured hdLECs, OSS led to the phosphorylation of eNOS coinciding with the phosphorylation of AKT. In contrast to blood vessel shear signaling, our data show that pharmacological inhibition of NO production does not have an effect on β-catenin activity. Similarly, we show that in hdLECS, *NOS3* knockdown led to reduced phospho-GSK3β, however, the pharmacological inhibition of NO production or PKG activation did not affect GSK3β phosphorylation. Thus, we show that eNOS regulates β-catenin activity independent of NO production. As an alternative explanation, we show that eNOS forms a complex with β-catenin and their binding is enhanced by exposure to OSS. Future studies are needed to address the cellular compartment in which this interaction takes place and whether disrupting binding can interfere with mechanotransduction signaling.

### eNOS regulates β-catenin nuclear translocation

In LECs, the adherens junction protein VE-cadherin binds β-catenin at the cell membrane^12^. During valve development, VE-cadherin releases β-catenin in response to OSS, thus allowing it to translocate to the nucleus and upregulate valve genes^12^. Additionally, the fraction of cellular β-catenin that can translocate to the nucleus is regulated by a multiprotein destruction complex involving GSK3β^21^. The complex sequentially phosphorylates β-catenin, targeting it for polyubiquitination and proteasomal degradation^21^. We found that in *Nos3^−/−^* embryos non-phosphorylated β-catenin was restricted to the membrane and absent in the nucleus. Consistent with this, we failed to detect nuclear β-catenin in *NOS3-*knockdown hdLECs exposed to either static or OSS flow conditions. We further showed that *NOS3-*knockdown hdLECs had increased GSK3β activity which led to lower levels of total β-catenin. Altogether, our data suggests the eNOS regulates β-catenin activity by preventing β-catenin degradation and promoting its nuclear translocation.

Since our data showed that eNOS regulates β-catenin activity, we attempted to rescue the valve defects in *Nos3^−/−^* mice by deleting exon 3 of the β-catenin gene which would prevent β-catenin from getting phosphorylated and degraded. Unexpectedly, we found that genetic overexpression of β-catenin in such a manner failed to increase the valve number in *Nos3^−/−^* mice at E18.5. It is possible that eNOS forms a complex with β-catenin to chaperone its nuclear translocation and would explain why preventing β-catenin degradation alone in the absence of eNOS would be insufficient to rescue valve formation in *Nos3^−/−^* embryos.

### Distinct signaling pathways regulate valve development during embryonic and postnatal stages

Unexpectedly, we found that at P7 there is no difference in lymphatic valves between WT and *Nos3^−/−^* mice, indicating that *Nos3^−/−^* mice experience a delay in valve development due to defective valve specification. Our data show that the percentage of branchpoints with valves increases in *Nos3^−/−^* mice from P3 to P7, indicating that *Nos3^−/−^* mice are actively growing new valves to make up for the initial delay in valve development. Since this rapid valve growth is independent of eNOS, we propose that other signaling pathways become active after birth to enable normal valve growth. Additionally, we show that postnatal loss of β-catenin only leads to a 55% loss of lymphatic valves whereas embryonic loss of β-catenin was reported to lead to a complete loss of valves^11^. This argues that a pathway independent of β-catenin is regulating postnatal valve development. Indeed, we recently published that FOXO1 is a target of OSS-induced AKT, so it may represent one alternative pathway regulating valve development postnatally^13^. FOXO1 acts as a repressor of valve genes such as FOXC2, GATA2, KLF4 and KLF2 and genetic deletion of *Foxo1* was found to promote new valve growth embryonically as well as postnatally^13^. Consistent with this, we were able to fully rescue the valve number in *Nos3^−/−^* embryos by genetically deleting *Foxo1*. The differences in signaling pathways that regulate lymphatic valve development in embryonic and postnatal mice could be governed by differences in lymph shear stress values between the two stages. The average lymph flow shear stress was measured to be 0.64 dynes/cm^2^ with peaks of 4-12 dynes/cm^2^ in adult rat mesenteric lymphatic vessels^24^. During embryonic development, a lower blood pressure would lead to a lower interstitial fluid load and thus a lower shear stress than that experienced postnatally after a rapid rise in blood pressure^25–27^. Our findings could imply that more FOXO1 is phosphorylated and excluded from the nucleus after birth to augment lymphatic valve development and explains the distinct phenotypes seen between embryonic and postnatal stages upon genetic deletion of β-catenin and explains how *Nos3^−/−^* mice are able grow new valves after birth. In conclusion, we provide a novel role for eNOS in regulating lymphatic valve specification and describe a mechanism by which eNOS forms a complex with β-catenin to regulate its transcriptional activity.

## ACKNOWLEDGMENTS

The authors thank Taija Makinen, Young-Kwon Hong and Makoto Taketo for kindly providing the *Prox1CreER^T2^, Prox1-GFP* and *Ctnnb1^lox(ex3)^* strains respectively.

## STUDY APPROVAL

All animal experiments were performed in accordance with the guidelines approved by the University of South Florida (USF) Institution of Animal Care and Use Committee (IACUC).

## AUTHOR CONTRIBUTIONS

DI performed experiments, analyzed data and wrote the manuscript. DM, KL and RB performed experiments. YY performed experiments, analyzed data and edited the manuscript. JPS analyzed data and wrote and edited the manuscript. All authors contributed to and approved the final version of the manuscript.

